# Phase composition-specific behaviour of functional RNAs in liquid-liquid phase-separated microenvironment

**DOI:** 10.64898/2026.05.19.726130

**Authors:** Aritra Chakraborty, Faizal Khan, Srishti Sharma, Sandeep Ameta

**Affiliations:** Department of Biology, Trivedi School of Biosciences, Ashoka University, Sonipat, India

**Keywords:** RNA-ligand interaction, phase diagram, phase-separated droplets

## Abstract

The internal dynamics of liquid-liquid phase-separated systems are governed primarily by polymer packing, excluded-volume effect, and interactions between polymers and encapsulated macro-molecules. Although one immediate effect of such a constrained microenvironment is diffusion limitation, it remains unclear whether encapsulated macromolecules can also exhibit phase composition-specific functional behaviour that is not observable in a well-mixed aqueous environment. In this regard, different phases in a phase-separated environment can be accessed via a phase diagram that demarcates the region between two-phase (droplets) and one-phase (polymer-rich, no droplets) regimes. While the two-phase region is heterogeneous, most previous work on encapsulating functional macromolecules in phase-separated droplets uses a single point from the phase diagram. This leaves a clear gap in understanding on how the function scales across this landscape of droplets and identifying regions advantageous for the encapsulated macromolecule and its function. Here, using the Spinach light-up RNA aptamer, we show that RNA function does not scale uniformly across the phase diagram. We show that RNA can exhibit phase composition-specific functional behaviour due to constraints imposed by the internal microenvironment of phase-separated droplets. Furthermore, using variants of the Spinach aptamer, we show that fluorescence activity differences among the variants vary differently with phase-separation regimes across the phase map, suggesting that some regions of the phase diagram can confer a selective advantage.

Our results highlight the potential of liquid-liquid phase-separated internal microenvironments in guiding the differentiation of functional RNA variants, which could serve as a physical selection pressure in pre-cellular evolution.

## Introduction

Despite extensive knowledge of the parameters that shape the functional fitness of structured RNAs, the impact of the complex internal microenvironment of liquid-liquid phase separation (LLPS) on the functional behaviour of RNA remains poorly understood. In biological systems, LLPS forms several membraneless subcellular compartments and plays a crucial role in the spatiotemporal control of biochemical processes (Banani et al. (2017); Hyman et al. (2014)). The malfunctioning of the LLPS process in cells has also been associated with numerous pathological conditions (Alberti and Dormann (2019); Ding et al. (2024)). In addition to their cellular functions, owing to their highly efficient partitioning of various functional biomolecules, LLPS droplets have also been envisaged as a compartmentalisation model (‘*protocell*’) that could have driven pre-cellular evolution (Haldane (1929); Oparin et al. (1957)). Supporting this, several studies have integrated different ‘*life-like*’ features, such as RNA catalysis (Strulson et al. (2012); Drobot et al. (2018); Poudyal et al. (2019a); Ameta et al. (2023), replication Poudyal et al. (2019b)) or self-reproduction of RNAs (Ameta et al. (2023)), metabolic activities (Smokers et al. (2022); Cao et al. (2025)), and growth-division scenarios (Zwicker et al. (2017); Ianeselli et al. (2022); Nakashima et al. (2021)). Nevertheless, the internal environment of these droplets is highly crowded and packed, which can significantly affect the dynamics of encapsulated biomolecules and, consequently, perturb their function. Furthermore, in charged systems (complex coacervates), intermolecular interactions can affect functional activity (Iglesias-Artola et al. (2022); Nakashima et al. (2025)), transport dynamics (Singh et al. (2024)), and duplex formation (Cakmak et al. (2020); Choi et al. (2022)). These interactions can be further exploited to modulate the material properties of LLPS droplets and to stabilise compartments (Le Vay et al. (2023)), thereby driving towards the coupling of protocellular activity to compartment growth and division (Zwicker et al. (2017); Sloot-beek et al. (2022)). The phase-separated compartment systems, in addition to supporting the function, provide several advantages to encapsulated biomolecules, *e*.*g*., preventing degradation (Okihana and Ponnamperuma (1982)) and rendering them robust to perturbations (Ameta et al. (2023)). Furthermore, recent work shows that, under temperature cycling, LLPS droplets can aid in distinguishing weak from efficient catalytic RNAs due to constrained diffusion at lower temperatures (Chakraborty et al. (2025)). Despite these advantages and their support for biomolecular function, it remains unclear whether the constrained microenvironment of LLPS droplets can enforce phase-composition-specific (or microenvironment-specific) functional behaviour that is not observable in a well-mixed aqueous system.

For biological systems, manifestation of environment-specific functions is well established, *e*.*g*., differential expression of the same gene in two different environments (Parts et al. (2012); Lappalainen et al. (2013); Mele et al. (2015)). In such a system, differential behaviour is often guided by a well-wired, complex, and highly regulated gene network comprising many components. However, it is not immediately clear whether a simple physicochemical system consisting solely of structured RNA, salts, and polymers can exhibit differential functional behaviour across two different environments. Furthermore, most studies analysing macromolecular function in the phase-separated environment capture only a few points on the full landscape of these ‘*liquid-like*’ droplets, creating a gap in understanding of how macromolecular functions scale across different areas (regimes) of the two-phase region from the phase diagram. Moreover, it remains unknown whether, when comparing two variants of the same RNA, there are any specific regimes in the phase diagram where the functional differences between closely related RNAs vary with the phase regime. As phase-separated droplets have also been envisioned as model protocell compartments and must support the evolution of encapsulated macromolecules, the ideal phase-separation regime is one in which the functional fitness differences among variants are maximised.

In this study, we demonstrate that the phase-separated microenvironment can drive phase composition-specific functional behaviour of RNA aptamers. Different phase-separation regimes lead to functional differences among sequence variants that are not apparent in a well-mixed aqueous environment (Fig. 1A). By comprehensively characterising several biophysical parameters across different LLPS regimes, we further show that this differential activity is dictated by the physical constraints imposed by the internal microenvironment of the phase-separated droplets.

**Figure 1.**
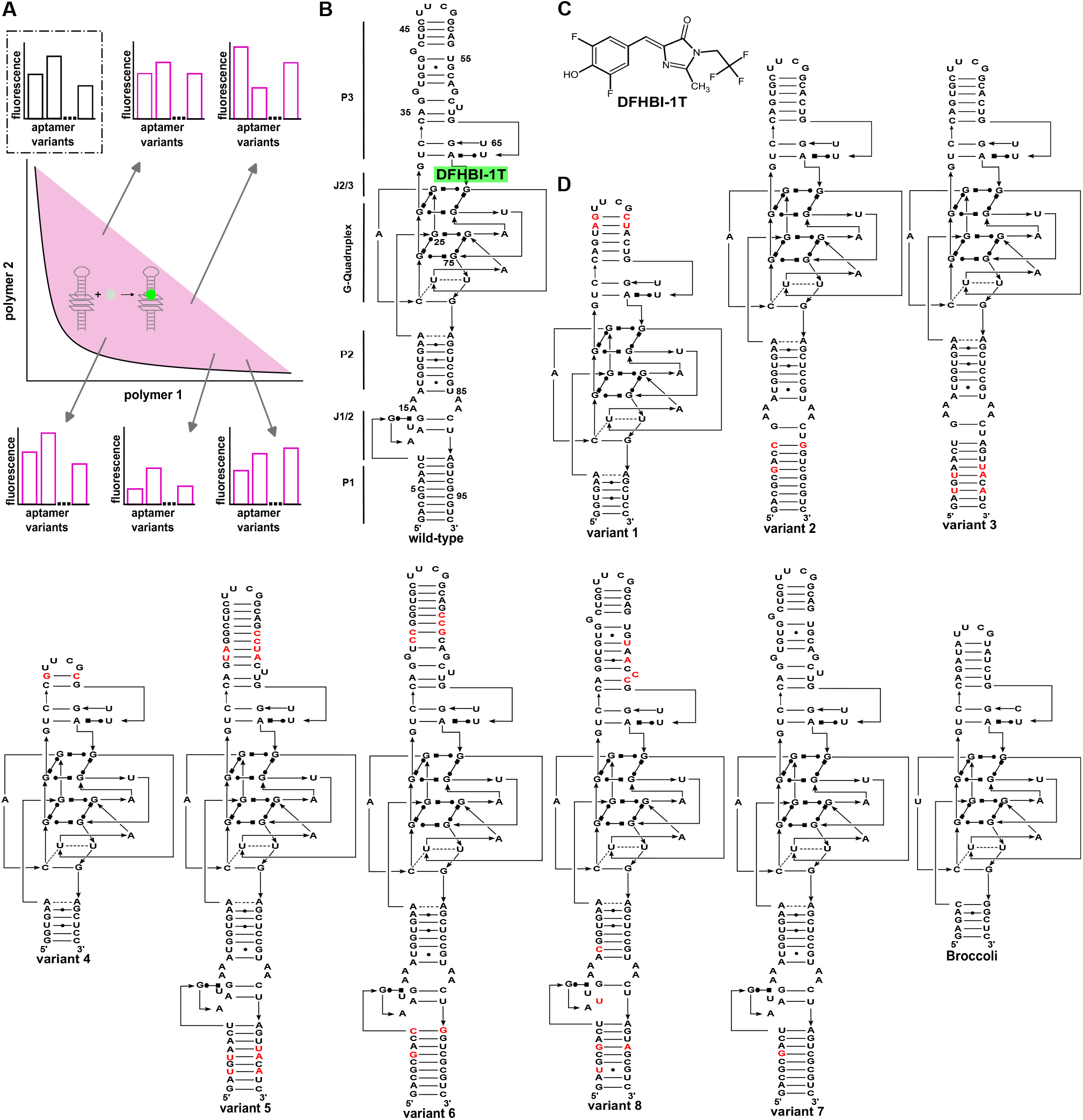
Phase composition-specific RNA function. (**A**). Schematic showing the scaling of the RNA function across the two-phase region (*magenta* area) of the phase diagram is non-uniform (histograms with *magenta* bars) and very different from what is observed in a well-mixed, non-phase-separated environment (*top left* histogram with *grey* bars). (**B**). The secondary structure of the Spinach RNA aptamer showing three paired regions (P1-P3), two connecting elements (J1/2, J2/3), and a central G-quadruplex with the upper two G-tetrads and a lower mixed tetrad. The ligand binds in the pocket formed by the upper G-tetrad and the U-A-U sequence stretch above it. (**C**). The structure of the ligand DFHBI-1T. (**D**). The predicted secondary structure of different ‘Spinach-like’ sequences (variants), including the Broccoli aptamer used in the study. These variants differ from the wild-type Spinach RNA aptamer sequence (Paige et al. (2011)) either by some nucleotide changes (highlighted in *red*) or deletion of sequence elements.

## Results

### Spinach RNA aptamer system and phase diagram

To investigate how the functional activity of RNA varies across the different regions of a phase diagram, we used Spinach RNA aptamer (Paige et al. (2011)) as a model system and measured its activity across the phase diagram (Fig. 1B). Spinach is an *in vitro* selected RNA aptamer against the green fluorescent protein (GFP) chromophore DFHBI ((Z)-4-(3,5-difluoro-4-hydroxybenzylidene)-1,2-dimethyl-1H-imidazol-5(4H)-one) (Paige et al. (2011)) which mimics the cyclic product of Ser65, Tyr66, and Gly67 in the GFP protein core (Ormo et al. (1996)). Although DFHBI exhibits minimal fluorescence on its own, its binding to Spinach RNA prevents *cis*-to-*trans* isomerisation across the ethylenic bridge, resulting in a GFP-like fluorescence enhancement over the background (Paige et al. (2011); You and Jaffrey (2015)), making it very useful for live-cell RNA imaging (Paige et al. (2012)). Several modifications to the DFHBI ligand have been explored to increase its photostability as well as *in vivo* applications (Song et al. (2014)). In the current study, we use DFHBI-1T, which has a trifluoroethyl group attached to the N1 of the imidazolone (Fig. 1C). The Spinach RNA aptamer has a coaxial structure with a conserved G-quadruplex (rG4) at its core, and the upper tetrad of this conserved rG4, along with a three-nucleotide stretch (U-A-U) above, constitutes the binding site for the DFHBI-1T (Fig. 1B, highlighted in *green*). Over the last few years, several ‘Spinach-like’ aptamers (Fig. 1D) have been thoroughly characterised with respect to their sequence composition (Okuda et al. (2017); Strack et al. (2013); Autour et al. (2016), structure (Warner et al. (2014)), and photophysics (Warner et al. (2013); Dao et al. (2021)).

For generating the phase-separated droplet landscape, we experimentally constructed a phase diagram by titrating different concentrations of polyethylene glycol (PEG) and dextran (Dex) (see Materials and Methods), resulting in a clear 2-dimensional landscape representing combinations of concentration where polymers segregate themselves from a one-phase region (clear solution) and turbid two-phase region, *i*.*e*., droplets and aqueous solution around them (Fig. 2A, *magenta* coloured area). To assess RNA functional activity, we selected five regions (referred to as regimes I to V; Fig. 2A, *right*) spanning the phase diagram that encompass a range of PEG-Dex stoichiometries. All these five regimes result in a turbid solution containing phase-separated droplets (Supplementary Fig. S1). The PEG-Dex system forms segregative phase-separated droplets, in which one polymer phase separates (dextran in this case), and the other is excluded (PEG in this case), and is also known as an aqueous two-phase system (Dignon et al. (2020); Martin (2019)). Using a fluorophore-doped dextran, we indeed confirm that irrespective of the PEG-Dex ratio, droplets from all five regimes are dextran-rich (Supplementary Fig. S2). Each of these five regimes was further characterised for its ability to encapsulate Spinach-ligand complexes, droplet size distribution, and droplet number density (Supplementary Fig. S3). Despite being internally dextran-rich droplets, each regime of the phase diagram was able to encapsulate the Spinach-ligand complexes, though with different efficiency (Fig. 2B). Furthermore, the mean droplet size varies between regimes (3.17 *µm* - 42.84 *µm*), with regime 4 being the smallest (Supplementary Fig. S3 A-E).

**Figure 2.**
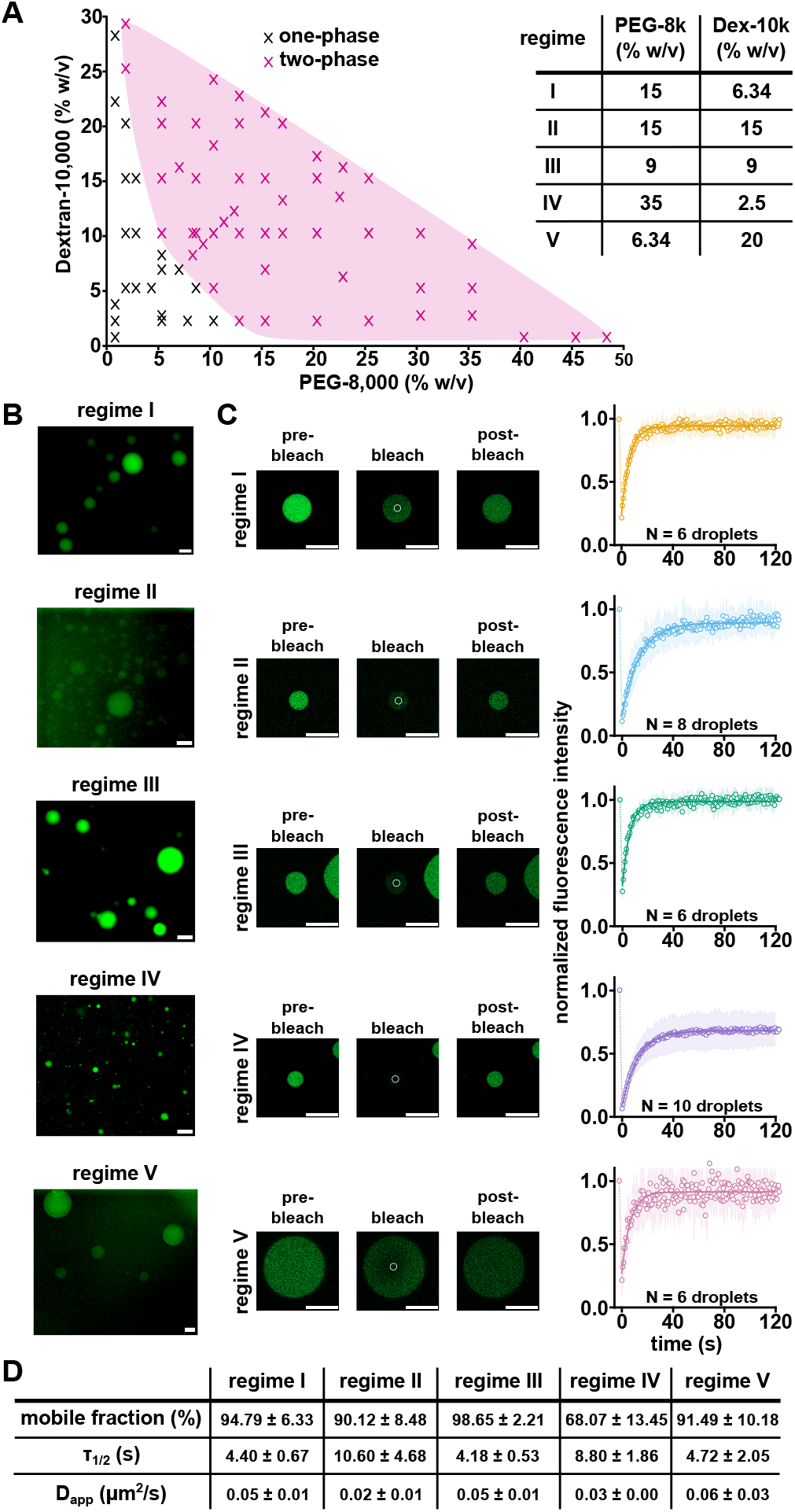
Characterisation of the phase-separated regimes. (**A**). Phase diagram of segregative phase separation exhibited by PEG-8,000 and Dex-10,000 (see Materials and Methods) showing a two-phase region (*magenta*) and one-phase region (*non-colored*). The cross marks represent the experimental data points. The polymer composition of different regimes is mentioned in the table in the *top right*. (**B**). Fluorescence microscopy images showing that the Spinach-DFHBI-1T complex remains functional (fluorescent) when encapsulated within phase-separated droplets across all regimes. The scale bar represents 15 *µm*. (**C**). The ‘*liquid-like*’ nature of the droplets is probed using FRAP performed on the droplets from all five regimes (see Materials and Methods). ***left*** ; Confocal fluorescence images of pre-bleach (*left*), bleached (*centre*) and post-bleach (*right*) droplets. All regimes show rapid fluorescence recovery post-bleaching due to the diffusion of the fluorescent Spinach-DFHBI-1T complexes from regions surrounding the bleach ROI (*white circles*). The scale bar represents 15 *µm*. ***right*** ; Fluorescence recovery curves extracted from the regimes. The solid lines represent the mean of at least 6 replicates, while the shaded regions around the solid line represent the standard deviation. (**D**). Table showing the quantification of the *mobile fraction*, the *τ*_1*/*2_, and the *D*_*app*_ of the fluorescence recovery across all the regimes.

We further characterise these regimes for ‘*liquid-like*’ behaviour and the transport properties of the encapsulated Spinach-ligand complexes by using FRAP (fluorescence recovery after photobleaching). All five regimes of the phase diagram were found to be forming ‘*liquid-like*’ droplets, as is evident by the fast fluorescence recovery, and the Spinach-ligand complexes were found to be mobile inside the droplets (Fig. 2C).

However, the regimes vary in the mobile fraction, recovery times, and apparent diffusion coefficients (Fig. 2D). The regime III has the highest recovery fraction (*∼*98 %) and fastest recovery time (*t*_1*/*2_ *∼*4 *sec*). Three other regimes (I, II, and V) have similar mobile fractions. Although regimes I, III, and V exhibit similar recovery times, regime II shows slower recovery (*t*_1*/*2_ =10.6 *sec*), despite being similar in the *mobile fraction*. Keeping the area of the bleached region constant, we compare the apparent diffusion coefficients, which vary only slightly across all five regimes. Because regime IV droplets are the smallest, the same bleach area across resulted in noisy FRAP measurements.

### Functional activity of RNA aptamer and its variants scales differently across the phase diagram

We next quantified the fluorescence of Spinach-ligand complexes in all five regimes of the phase diagram using a plate-based assay. While fluorescence increases significantly across all regimes relative to the ligand alone, the responses differ remarkably when compared to those without phase separation (Fig. 3A). A significant increase in fluorescence was observed in regimes I and V; fluorescence in regime II was also higher, whereas regime III was similar to what was observed without phase separation (Fig. 3A, *right, grey bar*). The regime IV showed less fluorescence. The regime IV droplets are also an order of magnitude smaller than other regimes. As a control, we also examined the fluorescence of the ligand alone, since an increase in local concentration within droplets could lead to unwanted fluorescence. However, ligand alone showed no in-crease in background fluorescence of DFHBI-1T (Fig. 3A, *left*). Similarly, another activity control, a sequence variant of the Spinach RNA aptamer, which was shown to lose the fluorescence activity (Okuda et al. (2017)), did not show any aberrant activity due to the crowded microenvironment of phase-separated droplets (Supplementary Fig. S4). These results confirm that RNA function varies across the phase diagram and it is not due to aberrant ligand-background fluorescence.

**Figure 3.**
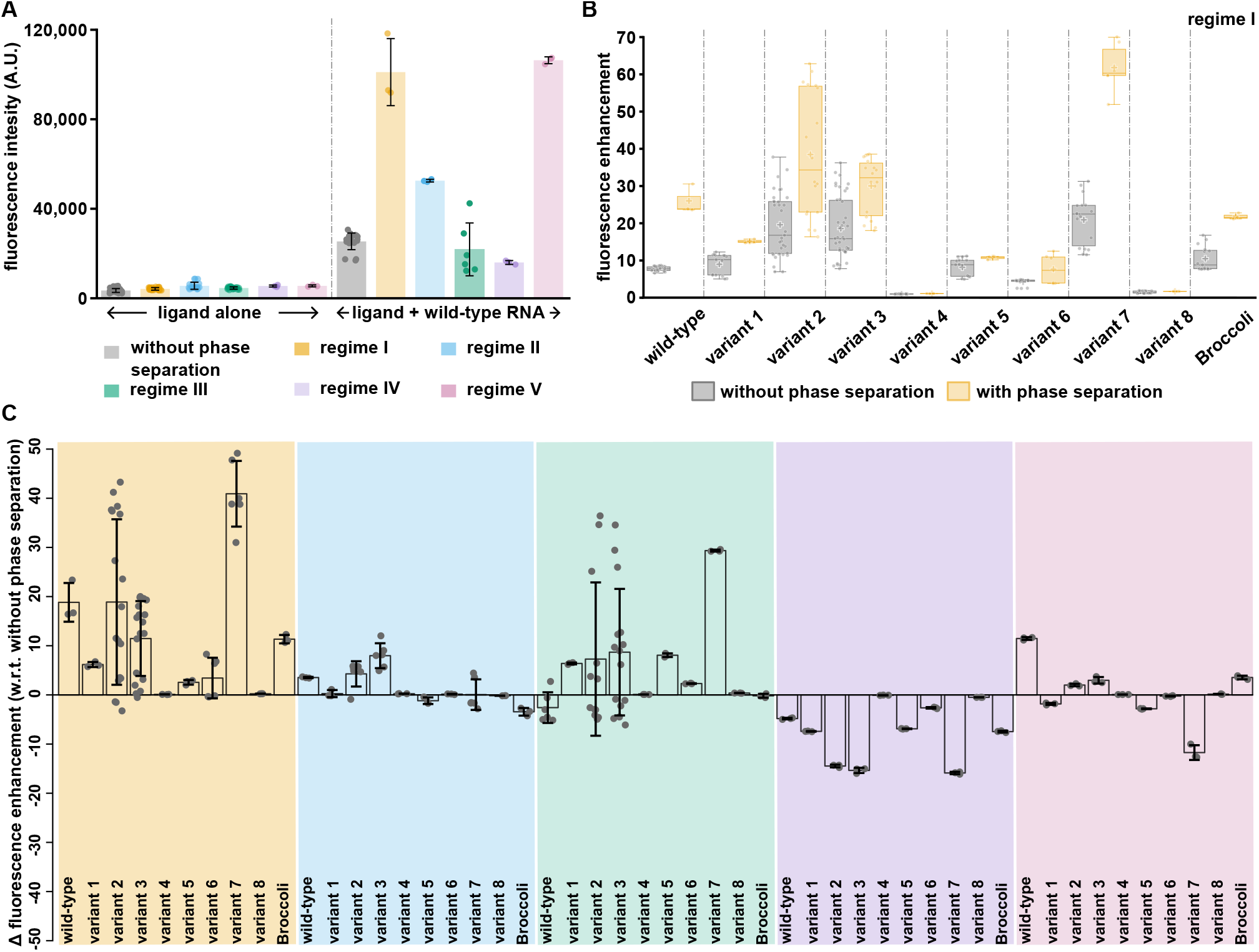
Functional output of the RNA variants scales differently across the phase diagram. (**A**). Bar plot showing the fluorescence intensity of wild-type Spinach RNA aptamer and ligand complexes (*right*) and ligand alone (*left*) across different phase-separated regimes. The individual data points are represented as overlaid solid circles. All measurements were performed in triplicate (or more), and the mean is presented with standard deviation. (**B**). Figure showing a box plot of the fluorescence enhancement (F_E_) of different RNA variants (‘Spinach-like’ sequences) in the phase-separation environment (regime I). The *grey* and *yellow* bars represent (F_E_) without and with phase separation, respectively. The boundaries of the box plots represent the first and third quartiles, while the lines within the boxes represent the median values; the whiskers represent the range of the data, and the plus/cross mark within the boxes represents the mean values. The individual data points are represented as overlaid solid circles. (**C**). Butterfly plot showing the ∆F_E_, the fluorescence enhancement (F_E_) of the RNA variants in the different phase-separated environments *w*.*r*.*t*. that without phase separation. The bars represent the mean values, with the error bars representing the standard deviations. The value of ‘0’ represents the same fluorescence enhancement as without phase separation, whereas values > ‘0’ or < ‘0’ show increased or decreased fluorescence enhancement in the phase-separated environment, respectively. The overlaid solid circles represent the individual data points.

While measurements with the wild-type Spinach RNA aptamer show functional variability across the phase diagram, from an evolutionary perspective, it is important to determine if phase-separation systems can aid in differentiating functional RNAs. To test this, we generated literature-reported (Okuda et al. (2017); Autour et al. (2016)) (Fig. 1D) as well as some rationally designed (Supplementary Fig. S5) ‘Spinach-like’ variants and measured fluorescence enhancement (F_E_: ratio of fluorescence intensity of the RNA-ligand complex to the ligand alone; see Materials and Methods) and compared it without phase separation. Interestingly, upon comparing RNA function in LLPS droplets to the well-mixed environment, there was an unexpected trend in their fluorescence activity, even within the same phase-separated regime (*e*.*g*., regime I, Fig. 3B, Supplementary Table S1). Several RNAs, which showed similar fluorescence intensity without phase separation (*e*.*g*., wild-type, variant 1, variant 5, and Broccoli), had a differential increase in fluorescence when they were subjected to a phase-separated environment, where wild-type increased significantly (F_E_ = *∼*26) whereas variant 5 gained slightly (F_E_ = *∼*10, Fig. 3B). Similarly, variant 7, which differs from the wild-type by a single-base mutation in the P1 region that increases fluorescence (Autour et al. (2016)), enhances significantly within the phase-separated environment (F_E_ = *∼*61). While other regimes also showed non-expected differential fluorescence enhancement (Supplementary Fig. S6), for a comprehensive overview, we plotted ∆ fluorescence enhancement (∆F_E_) of each RNA variant in the different phase-separated regimes *w*.*r*.*t*. that in the well-mixed, without phase separation environment (see Materials and Methods). When analysing ∆F_E_ across all the regimes, it becomes immediately apparent that the functional differences do not scale the same way in each regime (Fig. 3C). For example, in the regime I of phase-separated environment, all variants show more fluorescence than in the absence of phase separation (∆ *>* 0), whereas in other regimes only a few variants (*viz* variant 3, 5, and 7) are enhanced. Surprisingly, in regime V, apart from Broccoli and wild-type Spinach, all the other variants showed a decrease in fluorescence (Fig. 3C, Supplementary Fig. S6). Please note that Broccoli aptamer behaviour is similar to the wild-type; however, it has been selected via an independent *in vitro* selection but yielding the same conserved core rG4 structure (Filonov et al. (2014); Okuda et al. (2017)). The RNAs in regime IV also showed a decrease in fluorescence; it might also be attributed to a smaller droplet size compared to other regimes, however comparison within regime IV showed that not all RNAs are enhanced equally, and the trend is different compared to other regions of the phase diagram (Fig. 3C and Supplementary Fig. S6). Interestingly, wild-type and var7, which are same in length (98 nt), showed opposite behaviour when encapsulated in regime I and regime V (∆F_E_ < 0 for variant 7 in regime V). A similar behaviour was also observed for the variants 2 and 3 (80 nt each but with few mutations in the P1 region). Furthermore, the differences in ∆F_E_ also vary from one regime to another, *e*.*g*. differences between wild-type and variant 3 in regimes I, II, III, and V. We further confirmed the fluorescence differences observed here between wild-type and variant 7, as well as variants 2 and 3, using microscopy, where droplets containing these variant-ligand complexes were encapsulated and the trend was found to be respected there too (Supplementary Fig. S7).

These results indicate that ‘Spinach-like’ fluorogenic RNAs behave differently across different compositions of phase-separated environments, and the functional differences among the variants don’t scale the same way across the phase diagram, indicating that functional RNAs could exhibit phase composition-specific behaviour.

### Selective exclusion of ligand from dextran-rich droplets

The results above suggest phase separation can enable differentiation between functional RNA sequences. In a simplistic system comprising RNA, a ligand, and a buffer, the differential functional behaviour of RNA across different phase-separated regimes could be governed solely by physical parameters, including interactions with the polymer (dextran), ligand binding to RNA, partitioning of RNA and ligand, and diffusion. Therefore, we next systematically probed the parameters governing this behaviour. Polymers like PEG act as a crowder and impose an excluded-volume effect (Alfano et al. (2024)), whereas dextran interacts with RNA via electrstatic interactions and has been extensively used to assist RNA delivery in therapeutic interventions (Hu et al. (2021)); therefore, it is imperative to test the effect of these polymers on RNA function. However, our results show that the PEG and dextran alone do not have a similar effect on the fluorescence of RNA-ligand complexes (Supplementary Fig. S8), indicating that it is not crowding or interactions alone, but rather the microenvironment created by LLPS droplets, that dictates fluorescence enhancement and differences between the RNA variants.

Next, we tested whether the differential fluorescence activity is due to changes in the binding behaviour of RNA and the ligand within the phase-separated droplets. Ac-cordingly, we quantified binding efficiencies (*K*_*d*_; binding coefficients) by measuring dose curves at increasing concentrations of the DFHBI-1T ligand (see Materials and Methods). Surprisingly, the measurements showed that, in general, ligand binding to its cognate RNA is slightly weaker in the phase-separated regimes (Fig. 4). In the case of the wild-type, it has a *K*_*d*_ of *∼*0.68 *µM* without phase separation, which gets worse in all the regimes of phase separation, even though there is an increase in the fluorescence (*∼*1.08 *µM*, Fig. 4A, *top*). A similar result was found for variant 7, for which the *K*_*d*_ has increased inside the phase-separated environment (Fig. 4A, *bottom*). This suggests that when comparing with and without phase separation, the correlation between *K*_*d*_ and fluorescence enhancement is not respected. Furthermore, within the phase-separated environment, when comparing the wild type and variant 7, while the trends between fluorescence enhancements were respected *K*_*d*_ in regime I, it was not the case in the regimes III and V; *e*.*g*., in regime III, there is a significant difference in the fluorescence enhancement between the wild-type and variant 7 despite having a similar *K*_*d*_ (Figure 4, Supplementary Table S2). For other variants, similar trends were observed across the phase diagram. These anomalies suggest that the binding efficiency (*K*_*d*_) of the ligand to its cognate RNA in a phase-separated environment can be highly unpredictable and cannot alone explain the differential fluorescence behaviour observed.

**Figure 4.**
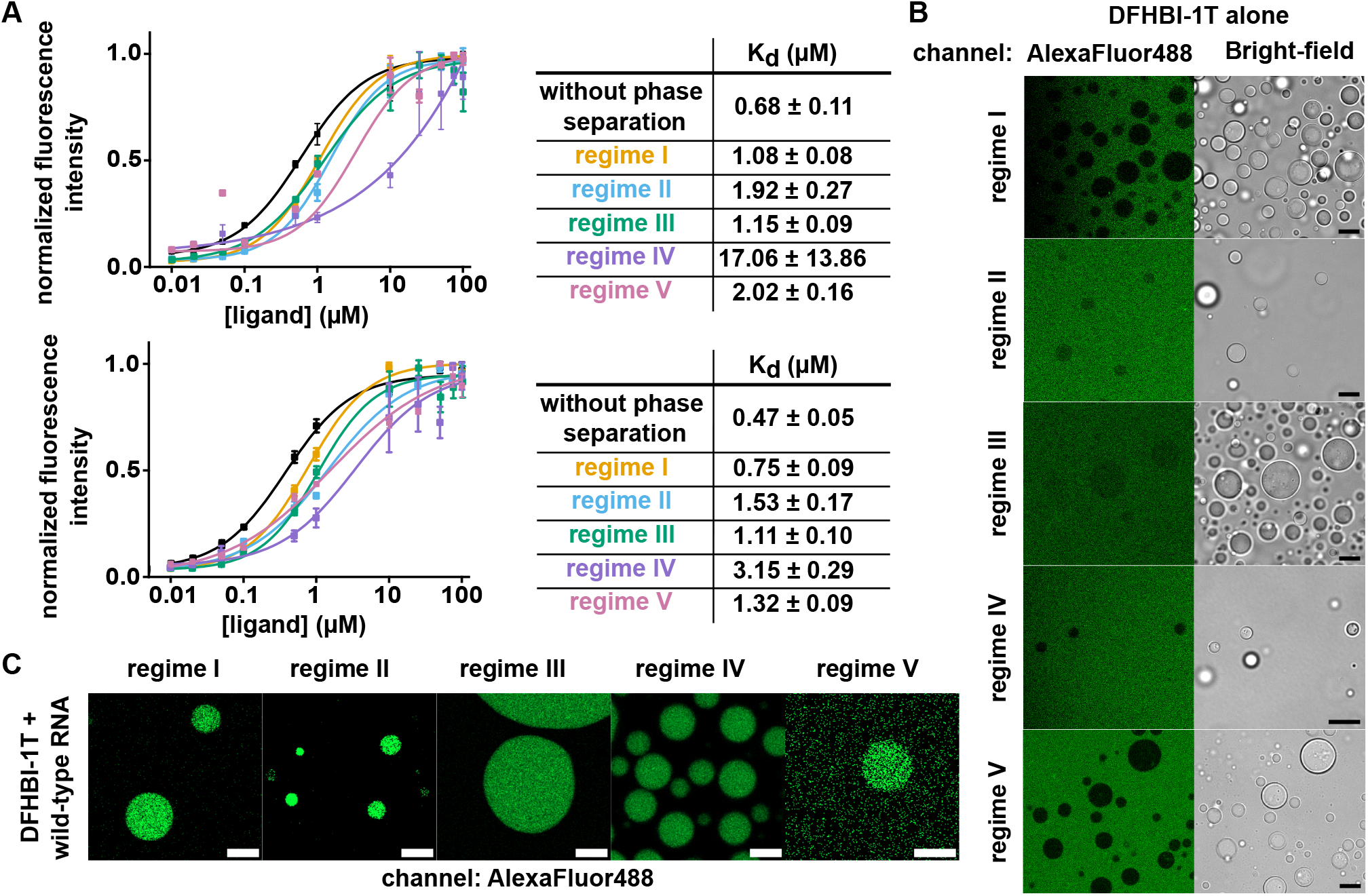
Different phase-separated environments affect the binding of the RNA-ligand complexes to different extents. (**A**). The *K*_*d*_ of wild-type RNA-ligand complex (*top*) and variant 7 RNA-ligand complex (*bottom*) in different environments. Each point represents the mean of three replicates (or more), with the error bars representing the standard deviation. The solid lines represent the Hill’s equation fit across the different environments. (**B**). The ligand, DFHBI-1T, does not enter the phase-separated droplets when present alone, as evident from the confocal images. Scale bar represents 15 *µm* in all regimes except in regime IV where it represents 5 *µm*. (**C**). In the presence of RNA (here, wild-type Spinach RNA aptamer), confocal images show that the RNA-ligand complexes enter the droplets, as indicated by higher fluorescence intensity within them. The scale bar represents 15 *µm* in regimes I, II, and III, 3 *µm* in regime IV, and 5 *µm* in regime V.

To understand this behaviour better, we further check the partitioning of RNA-ligand complexes and the DFHBI-1T ligand alone inside these dextran-rich droplets from each regime by measuring the partition coefficient (*K*_*P*_). While generally *K*_*P*_ are carried out by averaging all droplets after separating from the rest of the liquid, we measured *K*_*P*_ at the single-droplet level (see Materials and Methods). When measured *K*_*P*_ for RNA-ligand complexes, we indeed observed that they are enriched inside the phase-separated droplets (*K*_*P*_ >1), the absolute values were found not to be significantly different from each other (Supplementary Table S3). Surprisingly, when measuring *K*_*P*_ for the DFHBI-1T ligand alone, interestingly, we observed that it is selectively excluded from the dextran-rich droplets irrespective of the phase separation regime (Fig. 4 B). However, when bound to RNA, it is partitioned inside the dextran-rich droplets (Fig. 4 C). The dextran-rich droplet creates a hydrophilic interior environment and may exclude the hydrophobic DFHBI-1T ligand. As a control, we examine another hydrophobic dye with and without oligonucleotide conjugation (Supplementary Fig. S9). As expected, the dye alone does not partition into the dextran-rich droplets, but when conjugated to nucleic acid, it is enriched within them. These results strongly hint that the lower *K*_*d*_ observed inside the phase-separation environment may result from the fact that only RNA-ligand complexes in a 1:1 ratio are being partitioned inside, leading to driving the binding equilibrium to a more off-state inside the phase-separated droplets. However, because small molecules and oligonucleotides exhibit constrained transport dynamics within phase-separated droplets (Singh et al. (2024)), which might not scale the same way as RNA and the ligand, assessing RNA-ligand complex binding on and off is not trivial and requires further exploration. Nevertheless, the results presented in this study indicate that RNA function, especially RNA-ligand interactions, does not scale uniformly across the phase diagram and can aid in differentiating functional RNA variants. The selective exclusion of a ligand may lead to distinct RNA-ligand binding dynamics within the phase-separated microenvironment, which are not observed in a well-mixed system.

## Discussion and Conclusions

Spatio-temporal organisation of functional biomolecules in a compartmentalised system is one of the key steps in the emergence of a ‘*life-like*’ chemical system on early Earth or in the design of synthetic cells. In addition to just physically enclosing the functional macromolecules, recent studies have shown that the compartment’s internal microenvironment can modulate stability and functional activity (Choi et al. (2022); Le Vay et al. (2023); Chakraborty et al. (2025); Iglesias-Artola et al. (2022)). Despite this, it is not immediately clear whether the constrained microenvironment of the phase-separated droplets can dictate phase-composition-specific functional behaviour and whether it can differentiate between RNA species. Differentiating functional RNA would be a critical feature of a protocell to drive molecular evolution in the pre-cellular era on Earth.

In the current work, we explore the landscape of phase-separated droplets, as mimicked by the phase diagram, to address the fundamental question of whether RNA function and functional differences between RNA variants scale in the same way as in the well-mixed, nonphase-separated environment. Using a well-established aqueous two-phase system of PEG-dextran droplets, which provides a dextran-rich, hydrophilic yet non-charged droplet into which charged nucleic acid can be encapsulated, we generated different microenvironments. To probe functional properties, we chose a fluorogenic RNA aptamer system based on the *in vitro* selected Spinach aptamer (Paige et al. (2011)), in which fluorescence output can serve as a good proxy for RNA-ligand interaction. We further employed ‘Spinach-like’ sequences (Okuda et al. (2017); Autour et al. (2016); Paige et al. (2012, 2011)) as variants to probe the scaling of differences in the functional output of these fluorogenic aptamer variants across the phase diagram. We observe that the RNA function, fluorescence from the Spinach aptamer, is not uniform across the phase diagram, and differences among the variants scale differently in different microenvironments. Interestingly, these differences were not observable in a well-mixed environment without phase separation. Such a feature confers a ‘selection agent’ role to the phase-separated compartment in addition to encapsulating and enriching the functional biomolecules. The phase-separated droplets have been indicated earlier to provide a selective advantage to the encapsulated functional molecule, *e*.*g*., poor and good RNA self-assembly under thermal cycling (Chakraborty et al. (2025)) or segregating RNA over DNA in peptide coacervates (Nakashima et al. (2025)), but studies exploring the differentiating RNA species across the landscape of droplets are lacking.

Another interesting feature of the current system is the selective exclusion of the hydrophobic ligand (DFHBI-1T) from the dextran-rich droplets, which has a significant impact on the binding constants that is not observable without phase-separation. While phase-separated droplets have been shown to have differential partitioning of molecules based on their chemical nature (Thody et al. (2024); Kelley et al. (2025)), in our case, the ligand can only partition inside the dextran-rich droplets in the bound state to RNA. However, once inside the droplets, it becomes non-trivial to understand how the binding-unbinding of the ligand proceeds. Since the inside environment is constrained (Singh et al. (2024)) as well as ligand-excluded, the availability of the ligand becomes diffusion-limited. Although we observe an apparent impact on the measured binding constants, a future study will require comprehensive, sensitive measurements to understand the diffusion of small molecules in such a constrained microenvironment and its impact on the selective advantage of encapsulated biomolecules.

The current study highlights the importance of exploring the landscape of protocells to identify ‘Goldilocks’ regions where the functional efficiency, as well as differentiating capability of the internal microenvironment, which will be crucial to driving the molecular evolution in a primordial environment or developing synthetic ‘*life-like*’ mimics.

## Materials and Methods

### Materials

All chemicals were purchased from Sisco Research Laboratories (SRL Chemicals Pvt. Ltd., India) and Sigma Aldrich, unless specified otherwise. T7 RNA polymerase was purchased from New England Biolabs (Product no.: M0251L). DFHBI-1T was purchased from Sigma (Product no.: SML2687). SYBR− Safe DNA gel stain (Product no.: S33102) was purchased from ThermoFisher Scientific. For all the experiments, nuclease-free water was used from SRL Chemicals (Product no.: 96370) or prepared by treating double-distilled water with diethyl pyrocarbonate (DEPC) from SRL Chemicals (Product no.: 46791). For denaturing gel preparation and running, acrylamide MW = 71.08 *g/mol* (SRL Chemicals, Product. No.: 89314), N,N-Methylene bisacrylamide (bis-acrylamide) MW = 154.17 *g/mol* (SRL, Product. No.: 38516), urea, MW = 60.06 *g/mol* (SRL, Product. No.: 69120), ammonium persulphate, MW = 228.20 *g/mol* (SRL Chemicals, Prod. No.: 28575), and N, N, N, N-Tetramethyl Ethylenediamine (TEMED), MW = 116.21 *g/mol* (SRL Chemicals, Product. No.: 84666, tris hydroxymethyl aminomethane hydrochloride, (Tris HCl), MW = 157.60 *g/mol* (SRL Chemicals, Product no.: 99438), boric acid, MW = 61.83 *g/mol* (SRL Chemicals, Product No.: 80266), EDTA (ethylenediaminetetraacetic acid disodium salt dihydrate), MW = 372.24 *g/mol* (SRL Chemicals, Product no.: 43272), NaOH pellets (sodium hydroxide pellets, MW = 40 *g/mol* (SRL Chemicals, Product No.: 68151) was used.

The nucleotide sequence of all RNA and DNA used in the study are listed in the Supplementary Tables S4-7. DNA primers and template oligonucleotides were purchased from Sigma-Aldrich and Barcode Biosciences and are listed in Supplementary Table S6, S7.

## Methods

### Transcription of RNAs

All the RNAs (Supplementary Tables S4, S5) were prepared using an *in vitro* run-off transcription protocol similar to that mentioned earlier (Ameta et al. (2023)). Double-stranded DNA templates were PCR-amplified using custom-synthesised primers and templates in a BioRad thermocycler or Proflex 3*32 well PCR system (Thermo Fisher). PCR was performed by initial denaturation at 94°C for 5 minutes, followed by iterative 30-34 cycles (denaturing temperature 92°C for 1 *min*, annealing temperature 57°C for 1 *min* and extension 72°C for 1 *min*), followed by final extension at 72°C for 5 *min*. The PCR products were verified on a 2% agarose gel stained with SYBR™ Safe DNA gel stain. The amplified DNA was precipitated overnight or for 4 *hours* at -20°C using 60 *µL* of 3*M* sodium acetate (pH 5.2) and 1 *mL* of pre-chilled 100% isopropanol in a 1.5 *mL* tube. The pellet was separated by centrifugation at 16,800 *rcf* at 4°C for 60 *min*, dried in a vacuum desiccator, and resuspended in nuclease-free water and used directly for the transcription reactions. *In vitro* transcriptions of the PCR products were performed using 4 *mM* NTPs, 0.01 *mg/ml* BSA, 10 *mM* DTT, and T7 RNA polymerase 4 U/*µL* in 1*X* transcription buffer (40 *mM* Tris pH 8.0, 1 *mM* spermidine, 22 *mM* MgCl2, 0.01% Triton X-100), incubated at 37°C for 14 *hours*. RNA purification done with 12% denaturing PAGE containing 30% (*w/v*) acrylamide/bisacrylamide (19 : 1) solution, 8*M* urea, and run at 28 *W* with 1X TBE buffer pH 8.3 (68.5 *mM* tris hydroxymethyl aminomethane hydrochloride, 88.95 *mM* boric acid, 20 *mM* EDTA, pH 8.0 (ethylenediaminetetraacetic acid disodium salt dihydrate)). The gel was briefly visualised under UV light, the RNA band was excised, and the RNA was eluted by incubation at 37°C for 2 *hours* or at 20°C overnight in 550 *µL* of 0.3 *M* sodium acetate. The eluate was filtered through a 0.2 *µm* filter, and RNA was precipitated out with 60 *µL* of 3 *M* sodium acetate and 1 *mL* chilled 100% isopropanol at -20°C for 4 *hours* or overnight. The pellet was separated by centrifugation at 16,800 *rcf* at 4°C for 60 *min* and resuspended in nuclease-free water. RNA concentration was determined using a Nanodrop spectrophotometer (Thermo Fisher, NANODROP ONE), and purity was further confirmed by running a 12% denaturing polyacrylamide gel.

### Microscopy

To probe droplet formation and construct the phase map, 30 *µL* of each sample was loaded into well setups and analysed at 4X to 20X objective magnifications using the Olympus CX-23 microscope. The well setup was created by pasting a piece of double-sided tape (having a hole punched in its centre) on a glass slide, sealing it with nail polish on all sides, and covering it with a glass coverslip (Blue Star, HSN code: 70179010). To estimate the size distributions and number densities of droplets across the regimes, droplets encapsulating labelled DNA (5’-Alexa488-labelled, Supplementary Table S6) were used as cargo, loaded into the well-set-up system, and imaged with the Olympus BX-63 epifluorescence microscope. To perform the FRAP experiments, droplets encapsulating the wild-type Spinach RNA aptamer and ligand complexes were loaded in a sandwich setup (created by adding 20 *µL* of the sample on a glass slide and covering it with a coverslip) and imaged using the Zeiss CellDiscoverer7 (confocal imaging with LSM 910).

### Phase diagram and droplet preparation

A phase diagram was generated by mixing polyethylene glycol (PEG, average MW 8,000; Sigma, Product no. P2139) and dextran (Dex, average MW 10,000; Sigma, Product no. D9260) at different concentrations from a stock solu-tion of 50 % (w/v) PEG and 30 % (w/v) Dextran. For each concentration condition, a 50 *µL* reaction was prepared by adding the calculated volumes of each polymer for the respective point in the phase diagram, along with nuclease-free water. The mixtures were then thoroughly homogenised using vigorous vortexing (Digital vortex mixer Thermo Scientific, Model no.88882010) at 3000 rpm for 30 *sec*. Immediately afterwards, a small aliquot from each mixer was examined under a compound microscope equipped with brightfield optics (see *Microscopy* section above) to confirm droplet formation. Only after confirmation of droplet formation is a particular combination of polymer concentrations assigned a two-phase point in the phase diagram.

For the size distribution estimates, droplet samples were prepared by co-encapsulating a 30-nucleotide 5’-Alexa Fluor 488-labelled DNA oligonucleotide (custom-synthesised by IDT, Integrated DNA Technologies, Belgium; Supplementary Table S6) into the sample mix before polymer addition. The samples were added to the well setup (described above), and measurements were carried out on the Olympus BX-63 epifluorescence microscope using cellSens software. For each sample, at least three different fields-of-vision (FOVs) were imaged in the FITC channel. The fluorescence images were subsequently analysed using the Fiji software (Schindelin et al. (2012)). The images were segmented using manual thresholding and binarised with watershed segmentation to distinguish clumped droplets, if any. Subsequently, the maximum Feret diameter was measured for all the segmented droplets using the Analyze Particle tool with the circularity set to 1 to avoid measuring any irregular blobs. At least 100 droplets were analysed to estimate droplet size in each regime. The size distributions were plotted as histograms in OriginPro, with the dotted line indicating the mean diameter.

### Fluorescence Recovery After Photobleaching (FRAP) measurements

To probe the ‘*liquid-like*’ nature of the phase-separated droplets, we performed FRAP on all five regimes droplets. For FRAP, droplet samples were created (as described above) by encapsulating 1 *µM* wild-type Spinach RNA and 50 *µM* DFHBI-1T. A 50-fold excess of the ligand was used to ensure sufficient droplet fluorescence despite rapid photobleaching. The droplets were imaged at laser powers ranging from 0.2 % to 5 % across the regimes, keeping the laser power constant within the regime. The region-of-interest (ROI) sizes were kept at 2 *µm* across the regimes to ensure that the ROIs were placed well within the droplets, thereby measuring the impact of recovery kinetics inside the droplets, largely devoid of edge effects. Equal-sized circular ROIs were placed on separate droplets and outside the droplet to act as a reference readout (to account for photobleaching during image acquisition) and as a background readout (to account for noise), respectively. Five frames were captured pre-bleaching to maximise fluorescence intensity. All images were analysed on ImageJ (Schindelin et al. (2012)), data were processed, and regressions were performed on OriginPro. For analysis, in each frame of the droplet images, circular ROIs were drawn on two different droplets—one to bleach a droplet (*bl*) and another to account for photobleaching (*ref*) that occurs—and an equal-sized ROI was drawn to measure background fluorescence (*bg*). All ROIs were drawn sufficiently away from the droplet edges. Average intensities within the bleach, reference, on ImageJ (Schindelin et al. (2012)), data were processed, and regressions were performed on OriginPro. For analysis, in each frame of the droplet images, circular ROIs were drawn on two different droplets—one to bleach a droplet (*bl*) and another to account for photobleaching (*ref*) that occurs—and an equal-sized ROI was drawn to measure background fluorescence (*bg*). All ROIs were drawn sufficiently away from the droplet edges. Average intensities within the bleach, reference, and background ROIs for all frames were measured in ImageJ (Schindelin et al. (2012)), subsequently processed as reported earlier (Jia et al. (2014); Ameta et al. (2023)), and are shown below. The data was corrected for noise by subtracting the background values.

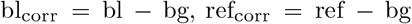

Photobleaching during image acquisition was accounted for as shown below.

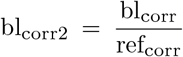

The photobleaching corrected values were normalized as shown below.

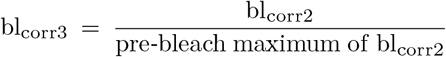

The normalised data were plotted on OriginPro to perform the regression. Only the post-bleach data was used for regression with a one-exponential decay curve.

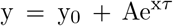

The offset *y*_0_ was used as a measure of the mobile fraction. The half-time of recovery was calculated as shown below.

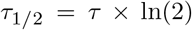

Using the Soumpasis model for a pure isotropic diffusion modelref, the apparent diffusion coefficients (*D*_*app*_) were calculated as shown below.

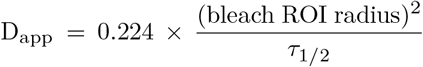

At least 6 droplets were analysed to estimate the *τ*_1*/*2_, the *mobilef raction*, and the *D*_*app*_ in each regime, and the values were reported as mean *±* standard deviation.

### Fluorescence measurements

To measure fluorescence enhancement upon ligand binding, the RNAs were first refolded by heating at 90°C for 2 *min*, then cooled to 4°C for 2 *min*. Following this, refolding buffer (40 *mM* Tris-HCl, pH 7.5, 5 *mM* MgCl2, 125 *mM* KCl) and DFHBI-1T (1 *µM*) were added, and the sample was incubated at 65°C for 5 *min*. The temperature was then gradually decreased by 1°C every 15 *sec* until it reached 25°C, and was maintained at 25°C for 15 *min*. Fluorescence measurements were performed using a CLARIO STAR plate reader in U-bottom 96-well plates at 25°C with the excitation wavelength of 455 *nm* and the emission wavelength of 506 *nm* (with a gain of 1400). Prior to each reading, the plate was shaken at 700 *rpm* for 60 *sec*. The fluorescence enhancement (F_E_) value of each RNA in each environment (phase-separated or well-mixed) was calculated by normalising to the average fluorescence of the ligand when present alone in the respective environment as follows:

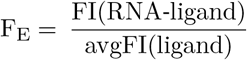

where, FI(RNA-ligand) is fluorescence intensity from the respective RNA variant along with the DHFBI-1T lig-and and avgFI(ligand) represents the average of triplicate measurements of the fluorescence intensity of the ligand alone. For the RNA-containing samples, at least 3 replicates were measured, and in some cases the number of replicates went up to 9. The data is reported as the mean *±*standard deviation. The ∆ fluorescence enhancement for each RNA in each environment was calculated as follows:

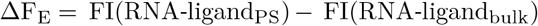

where, FI(RNA-ligand_PS_) is fluorescence intensity from the RNA-ligand complexes inside the phase-separated environment and FI(RNA-ligand_bulk_) represents fluorescence intensity from the RNA-ligand complexes without phase-separation.

### K_d_ measurements

The binding affinity of RNA toward DFHBI-1T was determined using fluorescence titration assays. RNA samples were prepared as previously described and mixed with different environments. The RNA concentration was kept constant at 0.2 *µM*, while DFHBI-1T was titrated over a range of concentrations typically from 0.01 *µM* -100 *µM*. Fluorescence measurements were recorded as described above. The fluorescence intensities measured for each RNA in each environment were normalised by taking ratios to the maximum fluorescence in the respective replicate. The normalised fluorescence intensities measured were plotted against the concentrations on a linear scale for analysis in OriginPro and fit using the Hill1 equation as follows:

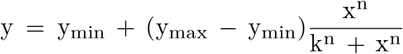

The *y*_max_ and the *y*_min_ represent the maximum and the minimum normalized fluorescence intensities, respectively, with the former being fixed at 1. The *k* and *n* represented the Michaelis constant and the number of cooperative sites, respectively. The value *K*_*d*_ was finally calculated *k*^*n*^ and reported as the mean *±*standard deviation from at least 3 replicates. For representation, the mean curves were plotted on a semi-log plot of normalized fluorescence intensity and concentration and fit using the same equation as above without keeping the *y*_max_ value fixed.

### Partition coefficient measurements

To ascertain the droplet partitioning coefficients (*K*_*p*_), droplet samples were prepared by adding 0.2 *µM* of *in vitro* transcribed RNA and 50 *µM* DFHBI-1T (ligand) to the sample mix before adding the polymers. The ligand was added in excess to account for photobleaching. The images were acquired using a method similar to that used for droplet size estimation. Following image acquisition, line regions of interest (LOIs) were drawn across droplets in each frame using the line selection tool in ImageJ (Schindelin et al. (2012)). Using the Plot Profile tool in ImageJ, fluorescence intensities across the LOIs were obtained and used to determine *K*_*p*_ as follows:

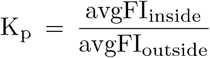

where, avgFI_inside_ is the average fluorescence intensity along *n* pixels inside the droplet and avgFI_outside_ represents the average fluorescence intensity along *n* pixels outside the droplet.

All images were acquired using the same exposure time within each regime. At least 50 droplets were analysed for each RNA and phase-separated regime combination, and the results were reported as the mean *±*standard deviation.

## Acknowledgments

We thank Manoj Kumar (Shiv Nadar University, Noida) and Philippe Nghe (ESPCI, Paris) for providing critical feedback on the manuscript and Shashi Thutupalli (National Centre for Biological Sciences, Bengaluru) for the constant support. We thank Sana Haan Jagmag and Ruchira Pal for their help with initial experiments. The authors thank Ashoka University for access to the common equipment facility. We also acknowledge access to the Ashoka-Zeiss microscopy and core microscopy facility at Trivedi School of Biosciences.

## Funding

S. Ameta acknowledge support from Anusandhan National Research Foundation Early Career Grant (ANRF/ECRG/2024/001288/LS) as well as Trivedi School of Biosciences, Ashoka University. A. Chakraborty and F. Khan acknowledge PhD fellowship support from Ashoka PhD program and Council of Scientific and Industrial Research, respectively.

## Author Contributions

S. Ameta conceived the idea and designed the study. A. Chakraborty and F. Khan performed all the experiments. S. Sharma helped in phase-diagram and K_*d*_ measurements. S. Ameta, A. Chakraborty and F. Khan interpreted the results and wrote the paper. All authors read and commented on the paper.

## Conflict of interest

None declared.

## Data Availability

The data that support the findings of this study are available from the corresponding authors upon reasonable request.

## Code Availability

Not applicable

## Notes

### Competing Interest Statement

The authors have declared no competing interest.

